# Lateral Undulation Aids Biological and Robotic Earthworm Anchoring and Locomotion

**DOI:** 10.1101/2021.02.02.429151

**Authors:** Yasemin Ozkan-Aydin, Bangyuan Liu, Alexandra Carruthers Ferrero, Max Seidel, Frank L. Hammond, Daniel I. Goldman

## Abstract

Earthworms (*Lumbricus terrestris*) are characterized by soft, highly flexible and extensible bodies, and are capable of locomoting in most terrestrial environments. Previous studies of earthworm movement have focused on the use of retrograde peristaltic gaits in which controlled contraction of longitudinal and circular muscles results in waves of shortening/thickening and thinning/lengthening of the hydrostatic skeleton. These waves can propel the animal across ground as well as into soil. However, worms can also benefit from axial body bends during locomotion. Such lateral undulation dynamics can aid locomotor function via hooking/anchoring (to provide propulsion), modify travel orientation (to avoid obstacles and generate turns) and even generate snake-like undulatory locomotion in environments where peristaltic locomotion results in poor performance. To the best of our knowledge, the important aspects of locomotion associated with the lateral undulation of an earthworm body are yet to be systematically investigated. In this study, we observed that within confined environments, the worm uses lateral undulation to anchor its body to the walls of their burrows and tip (nose) bending to search the environment. This relatively simple locomotion strategy drastically improved the performance of our soft bodied robophysical model of the earthworm both in a confined (in an acrylic tube) and above-ground heterogeneous environment (rigid pegs), where the peristaltic gait often fails. In summary, lateral undulation facilitates the mobility of earthworm locomotion in diverse environments and can play an important role in the creation of low cost soft robotic devices capable of traversing a variety of environments.

## 1. Introduction

Terrestrial animals have evolved strategies for effective propulsion and lift in complex terrestrial environments using a diversity of structures [1, 2], including whole bodies [3–5], heads [6] and limbs [7–10]. For example, several animal groups travel through/on flowable substrates via undulatory propulsion that involves the motion of the whole body [5, 6, 11, 12].

While many terrestrial locomotion studies focus on ground with rigid, flat, or homogeneous characteristics (e.g. dry sand) [13, 14], movement in cohesive substrates is less explored [15–17]. Among terrestrial animals, earthworms provide an excellent experimental system to discover principles of terrestrial locomotion in cohesive soils [1, 18, 19] as well as diverse terradynamic regimes. Earthworms locomote within cohesive soils for many different reasons, such as nesting, foraging, and surviving extreme environmental conditions (e.g. temperature, dryness etc.) [18, 20]. They create tunnels of varying size and shape through soil burrowing and organic matter burial behaviors [21–24]. They locomote in confined spaces and underground tunnels largely via retrograde peristaltic gaits where a transient substrate anchor is formed by the contraction of longitudinal muscles [1, 18].

The peristaltic gait that earthworms use to crawl within and on soil has been studied by many researchers [25–31]. These studies primarily focused on peristaltic crawling in a straight line, however, in reality, the environment in which the worms move has complex structures (e.g. different size and orientation tunnels, Fig. 2), requiring the worms to use other types of movement. In addition to crawling in a straight line, the worms change the shape of their long, slender bodies by bending or buckling to move on most surfaces, even on smooth, hard ground.

**Figure 1:**
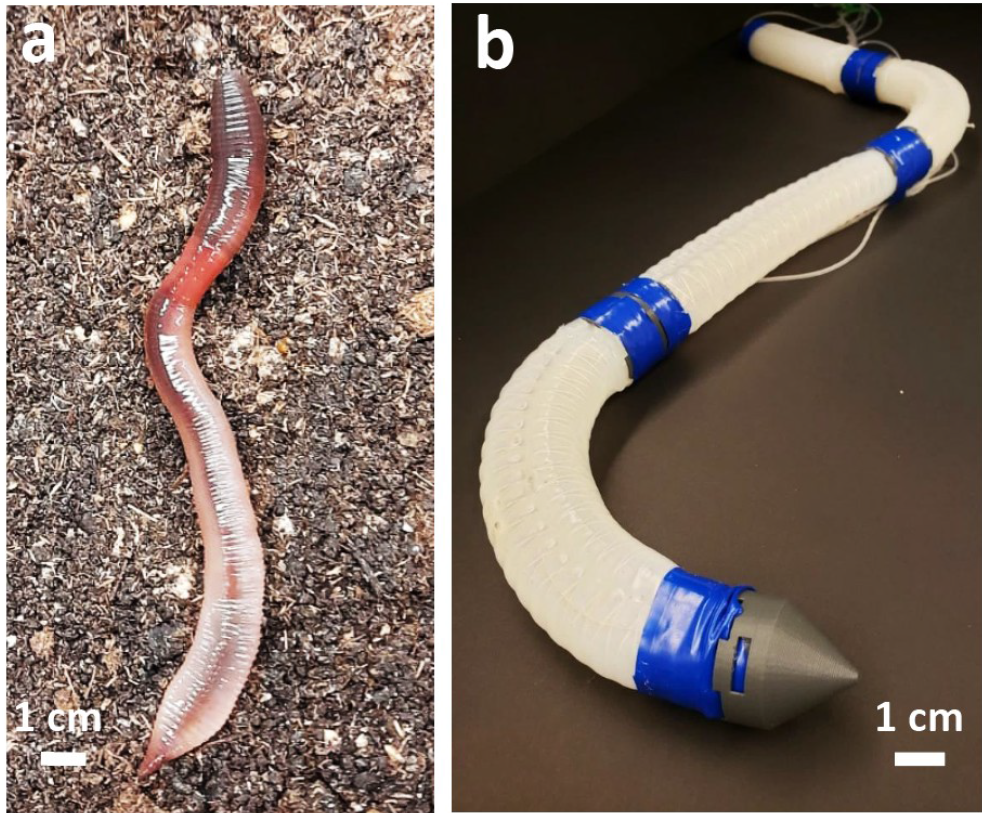
A biological earthworm and a robophysical model. (a) An earthworm (*Lumbricus terrestris*) on garden soil. (b) Fiber-reinforced, pneumatically driven segmented soft worm robot.

**Figure 2:**
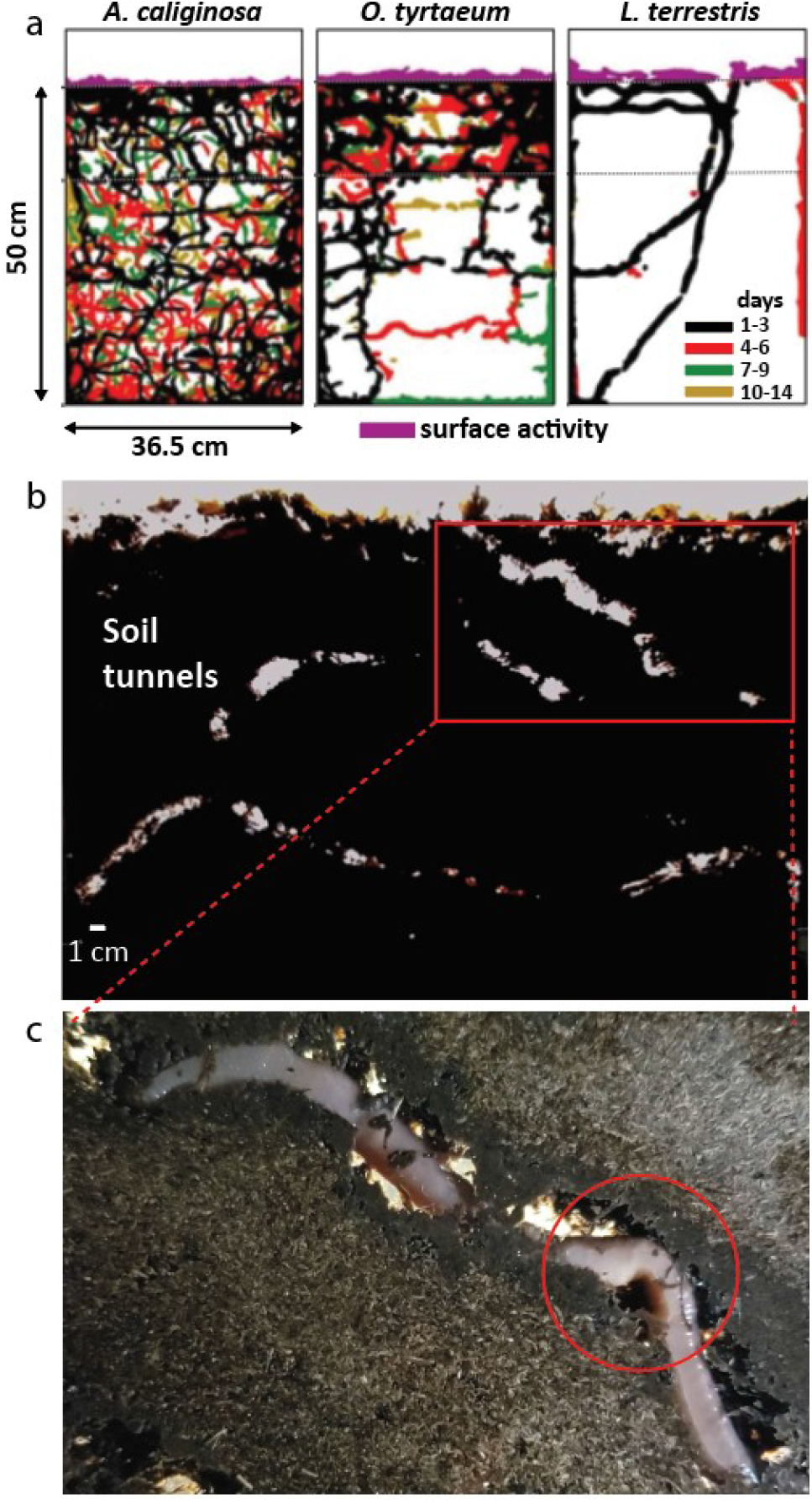
Earthworm tunnel dynamics and lateral undulation in a confined environment. (a) Burrowing patterns of three earthworm species (*A. caliginosa, O. tyrtaeum* and *L. terrestris*) in 2D terraria. The colors (black, red, green and gold) represent the days that the tunnels created (in total 14 days). Adapted from [21] with permission. (b) We monitored the activity of three earthworms in a 2D box (30×30×1.5 cm^3^) filled with moist garden soil. The figure shows the tunnels created by the worms during two weeks under 24*°*C room temperature. (c) The bending of the body (see red circle) during climbing in a large under soil tunnel.

The subterranean excavation ability of earthworms has inspired researchers to design earthworm-type robots [32–41], however, none of these robots burrow successfully in soil due in large part to high penetration forces and energy requirements [42,43]. Instead of burrowing, these robots locomote in structured tunnels and on the surface by mimicking the peristaltic motion of the worm, where the segments can either elongate longitudinally or expand radially to form anchor points. This approach of locomotion is useful if the radially extending segments can expand to fit the tunnel size; however, in most cases the segments cannot expand sufficiently due to the limited radial deformation. Moreover, being limited to pure peristalsis, robots cannot change their direction of movement, which reduces their mobility and maneuverability, especially in a heterogeneous environment.

Here, we discover that earthworms use lateral undulation when moving in models of their natural environments. Motivated by the locomotion capabilities of the worm, we design a robophysical model of an earthworm whose soft elastomer segments can elongate longitudinally (for peristaltic motion) and also undulate laterally in 2D space (for steering and anchoring). We systematically study how the earthworms control their shapes depending on the environment in which they locomote. Similar to the biological earthworm, we study the performance of our robophysical model in both confined and heterogeneous environments.

## 2. Methods and Results

Several studies were performed to investigate the burrowing behavior of earthworms under soil [21, 24, 44]. Depending on the earthworm species, worms create a network of horizontal and vertical tunnels in various sizes and orientations and use these tunnels for shelter and protection [21, 45, 46] (Fig. 2a).

Here, we particularly focus on how the worms locomote in their burrows using body undulation. In all experiments, we used deep burrowing anecic species (*Lumbricus terrestris*) of earthworms [47]. Before systematically examining the benefits of worms’ undulatory behavior, we first performed experiments in models of their natural environment, i.e. under soil tunnels, using 2D terraria described in [48]. Each terrarium was built by two parallel sheets of glass (30×30 cm^2^) and separated with 1 cm aluminum plates. We filled the terraria homogeneously with wet Magic Worm Bedding (moisture 80%) to construct a dark and moist model of their natural habitat.

Earthworms (n=10, L=26.95±6.18 cm, m= 7.5±2.04 g) obtained from Carolina Biological Supply are maintained in Magic Worm Bedding (Magic Products Inc., WI, USA) and kept in an environmental chamber (15*°*C, a relative humidity of 30%). We placed three earthworms in each terrarium and covered the top of the boxes with aluminum foil to prevent the worms from escaping. We kept the boxes in the environmental chamber one week prior to the experiments to make sure that the worms adapt to the experimental soil. After an adaptation week, we recorded the burrow patterns and activity of the worms in room temperature (24*°*C) for a week. To stimulate the movement of the worms in their tunnels, we put organic fruit particles at the top of the box.

Fig. 2b shows tunnels of the worms built in a week (see SI Movie-1 for timelapse video). We discovered that the worms employ lateral undulation and buckling of their flexible body segments to locomote in the tunnels. To quantify the occurrence of lateral undulation, we counted the frames that we observed body undulation (85 frames) during the first two days of the experiment (total 5760 frames) by focusing on the tunnel areas where the worms movement are clearly visible. We also recorded realtime video of the worms while moving in the tunnel. Fig. 2c shows a snapshot from the experiments where the worm used lateral undulation to anchor its body to the walls of the wide tunnel (SI Movie-1).

### 2.1. Systematic animal experiments in confined space (undulation aided anchoring)

In this section, we seek to understand how earthworms change their locomotion strategies in tunnels depending on the tunnel size and orientation. As we pointed out in the previous section, earthworms dig underground tunnels and spend most of their time moving inside these tunnels. Depending on the type of earthworms and environmental conditions, these tunnels can reach up to a meter [49]. To the best of our knowledge no one has systematically studied how worms adapt their gaits, especially how they use body undulation, to locomote in different size and angle tunnels.

In the experiments, we allowed earthworms to crawl in two different size acrylic tubes (*d*_1_ = 1 cm, *d*_2_ = 1.2 cm, length = 1 m, *d*_1_*, d*_2_ *>* worm diameter) while varying the tube angle from 0 to 90*°* with an increment of 15*°*. We calculated the average body length per gait cycle (BL/cyc) over several gait cycle which is defined with a body wave which starts at the anterior end of the body and moves towards the tail.

In each tube angle, we did at least one experiment with four different animals. To induce movement of the worms in the tubes, we stimulated the animals from their tail using a pointed object. Some of the animals were not able to crawl even with stimulation. We have not included these experiments in our results (in a tube *d*_1_=1 cm, 8 of 42 trials and *d*_2_=1.2 cm, 5 of 30 trials).

At a 0*°* incline, in both tube sizes the worms had the largest BL/cyc (for *d*_1_: 0.20 ± 0.075 and *d*_2_: 0.11 ± 0.015) and it had the lowest BL/cyc for *d*_1_: 0.076 ± 0.016 at a 90*°* incline and *d*_2_: 0.04 ± 0.018 at a 75*°* incline but could still climb without significant slipping (Fig. 3e). We measured the shape changes of worms while climbing in a 0 and 90*°* inclined tube. Fig. 4 shows the percentage of the body on the left and right side of the mid-line during locomotion. On level ground, the worms mostly used peristaltic gait by extending and contracting (shortened) its body in a cyclic way (Fig. 3-a,b). The shape of the body did not significantly change from cycle to cycle (Fig. 4b). At higher degree angles, such as 75 and 90*°*, body bending was more prevalent than at lower angles, providing extra anchoring points for forward locomotion and reducing the occurrences of slipping (Fig. 3-c,d). When they crawled in a 90*°* tube, they attempted to keep the body length constant and undulate their body to balance out the body forces (Fig. 4c). In the larger diameter tube (*d*_2_=1.2 cm) worms could not climb at 90*°*, however, they could maintain their position in the tube without slipping by buckling their body parts (Fig. 3f, SI Movie-2).

**Figure 3:**
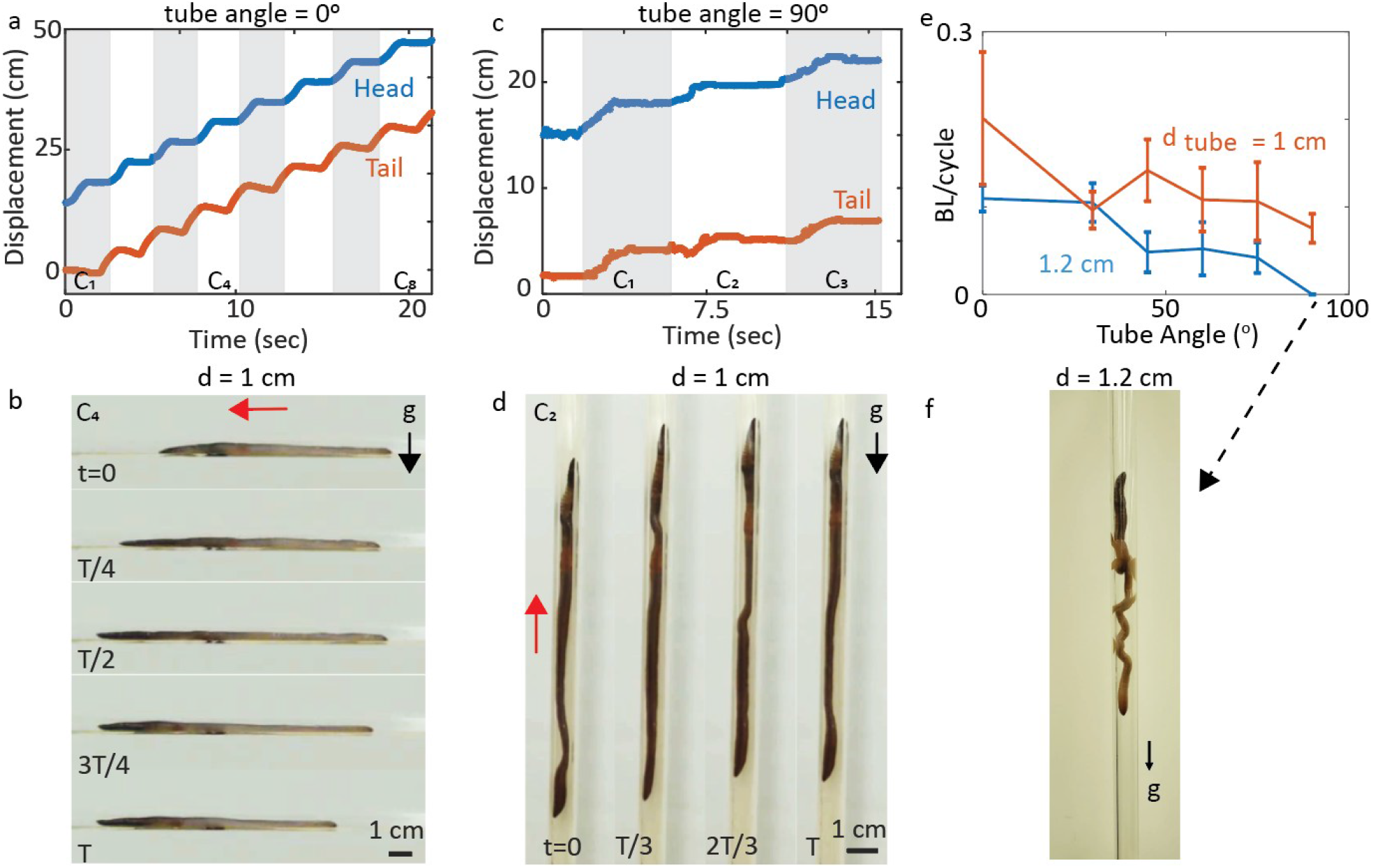
Lateral undulation in a laboratory confined environment. **(a)** Head (blue) and tail (red) trajectories of an earthworm when crawling in a 0*°* acrylic tube (*d_tube_* = 1 cm, the diameter of the tube is slightly larger than the worm diameter). White and gray areas show the each gait cycles (C_1_ to C_8_, total 8 cycles). **(b)** Example gait cycle (C_4_) from the experiment given in (a). Red arrow shows the direction of the locomotion. T represents the total time required for a one cycle. **(c)** Head (blue) and tail (red) trajectories of an earthworm when crawling in a 90*°* acrylic tube (*d_tube_* = 1 cm). White and gray areas show the each gait cycles (C_1_ to C_3_, total 3 cycles). **(d)** Example gait cycle (C_4_) from the experiment given in (c). Red arrow shows the direction of the locomotion and black arrow shows the gravity. **(e)** Displacement per gait cycle for tubes *d_tube_* = 1 and 1.2 cm as a function of tube angle *α* = 0, 30, 45, 60, 75 and 90*°*. **(f)** The worm prevents from sliding down by increasing the number of anchoring points on the body in a 90*°* acrylic tube (*d_tube_* = 1.2 cm).

**Figure 4:**
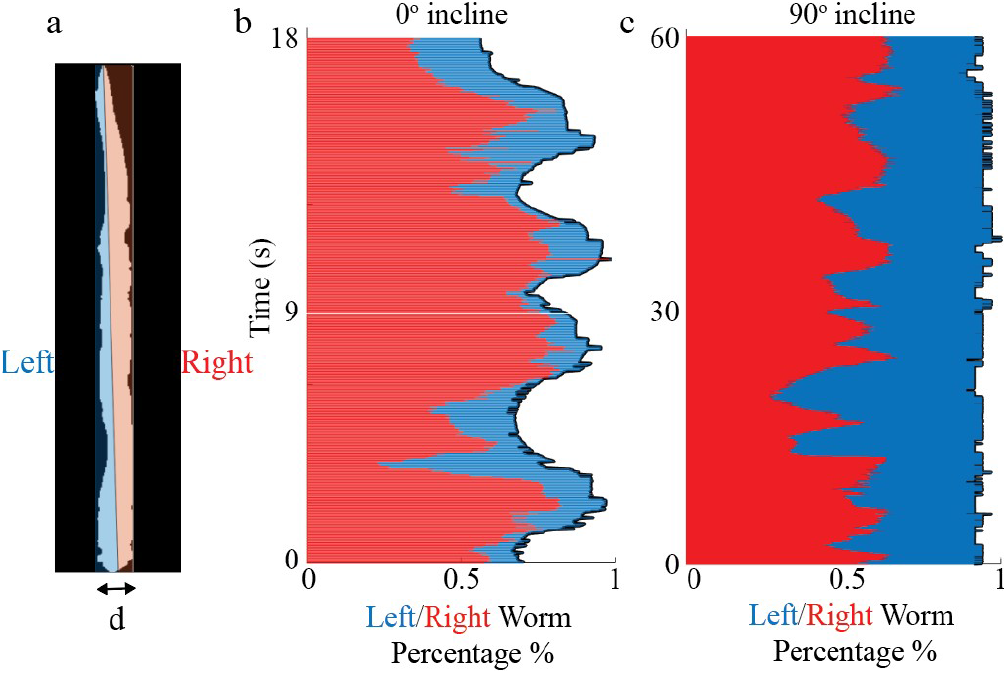
Worm shape changes during locomotion in a confined environment. **(a)** Colors (red:right, blue:left) show the area of the worm with respect to body mid-line (the line connects the head to tail). **(b)** Normalized left and right area of the worm when crawling in 0*°* and **(c)** 90 *°* inclined tube (d = 1 cm).

### 2.2. Confined space robot experiments

The animal experiments in various environments have revealed that successful locomotion of earthworms depends on the execution of lateral undulation which generate appropriate reaction forces from the environment. Our observations of lateral bending from the animal experiments given in previous sections encouraged us to develop a robophysical model of an earthworm whose novel mechanical design enables locomotion capabilities analogous to those of earthworms. The robot has a soft, flexible segmented body which can bend and extend with pneumatic actuation. In this section, we test our biological observation through robophysical experiments; in confined and heterogeneous environments.

#### Soft robot fabrication

We developed a worm robot by connecting four fiber-reinforced actuators in series (see Fig. 5a) using the methodology as described in [50]. Basically, fabrication involves multi-step molding of elastomer materials along with cylindrical geometry. Each actuator consisted of two half-cylinder elastomeric (Dragon Skin 10, Smooth-On) inner chambers that have helical fibers (Kevlar thread) wound around the outside (see Fig. 5a for the details of the fabrication steps). These two inner chambers allow the segment expand in directions with the lowest stiffness; i.e. the segment bends left/right when the one of the chamber is inflated and extend longitudinally when both of them are inflated (see right panel of Fig. 5b). Different than the previous pneumatically driven soft elastomer earthworm robots where the anchoring provided by radially expanding segments [35–38,41,51], here we use undulation of the segment to anchor and steer the robot in complex environments.

**Figure 5:**
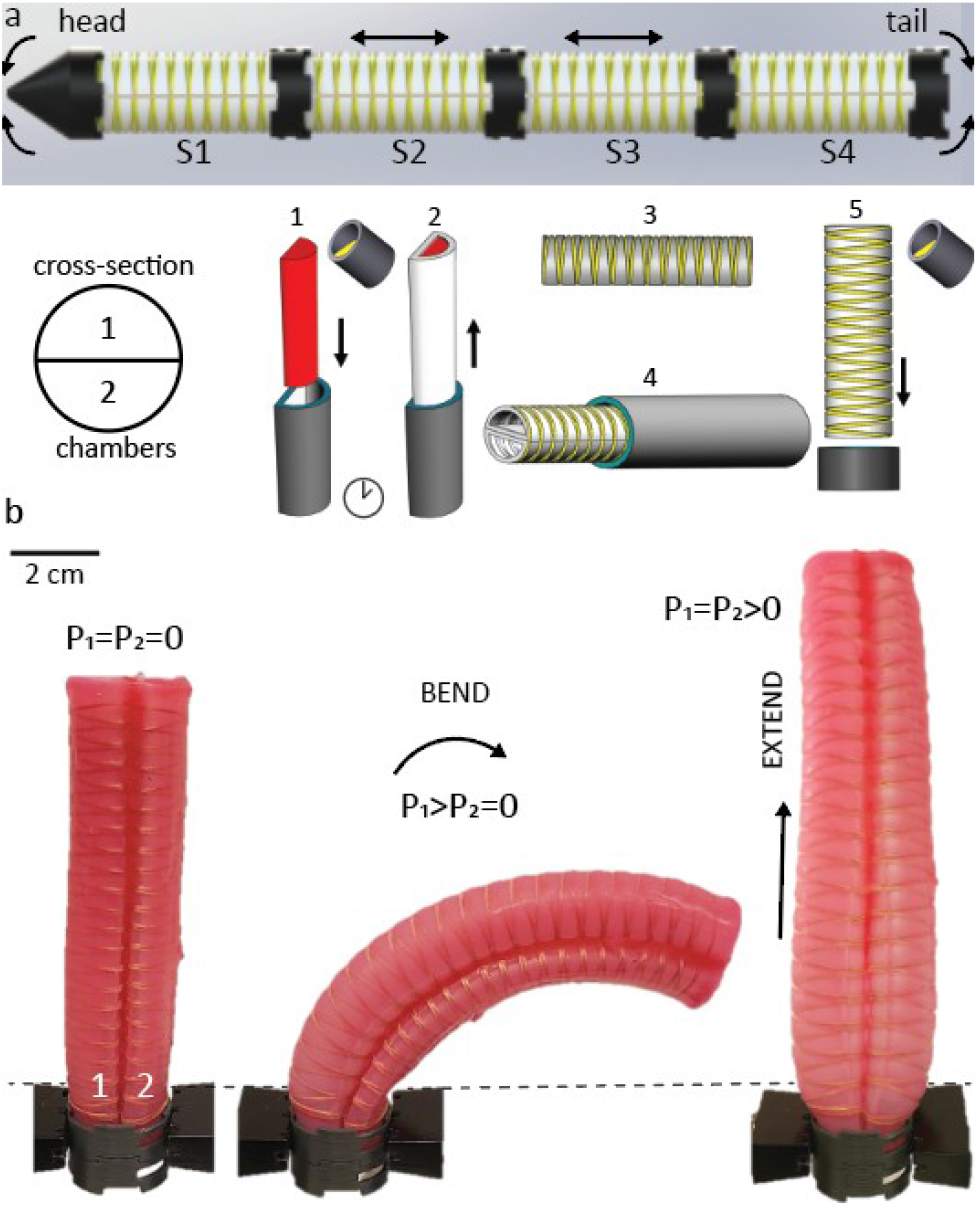
Design of the earthworm robot platform. (a) CAD diagram of the four segmented robot with fiber-reinforced elastomeric actuators. Each segments have two inner chambers with a radius of 1.25 cm, wall thickness of 2 mm, and length of 100 mm. The robot is 45 cm in total length, 2.5 cm in diameter. Arrows show the actuation type of the segments (straight: extension, curved: bending). Pneumatic tubes are routed outside of the body. Inset shows the fabrication steps (1-2) Molding and demolding of first layer. Uncured silicone (Dragon Skin 10) was poured into a 3D printed mold and was cured at 60*°*C for 4 hours. (3) Wrapping first layer with inextensible fiber (Kevlar thread, d=0.635 mm, Mcmaster) with a fiber angle 15*°*, (4) Attaching two inner chambers bilaterally to each other with a second layer of elastomer, (5) Molding silicone cap. (b) Example actuation states; (left) unpressurized state, the pressure of the both chambers are equal to zero (*P*_1_ = *P*_2_ = 0), (middle) bending state, pressure of the one of the chamber is bigger than other (*P*_1_ *> P*_2_ = 0). Inflated chamber produces a net curvature toward the uninflated chamber, (right) extension state, pressure of the both of the chambers are equal and greater than zero (*P*_1_ = *P*_2_ *>* 0),

The robot was assembled by attaching four actuators with 3D printed rigid connectors. We inserted straight barbed fittings (1/16 ID, McMaster-Carr) extended with Tygon PVC tubing (durometer 65A, 1/8 ID, 1/4 OD, McMaster-Carr) to the end of the actuators (at the side of connectors) and sealed it with Sil-Poxy glue (Smooth-On) for an air input. The silicone rubber tubings (durometer 50A, 1/32 ID, 1/16 OD, McMaster-Carr) were used to connect the actuators to an air source.

#### Control board

The robot was controlled with a setup consisting of an Arduino Mega 2560 microcontroller, eight solenoid valves (NITRA, 5-port, 4-way, 3-position, AutomationDirect), eight manual pneumatic regulators (Nitra, 4-57 adjustable range, Automation Direct), and IRF540 MOSFET switch modules. 4-way, 3-position solenoid valves allow the chambers to inflate, deflate, or hold the air pressure according to activation pattern. We control the pressure of the each chamber independently by using a network of pneumatic channels and Arduino-controlled solenoid valve for each segment. Each of the four segments could be pressurized from an external source (compressed air, 10-15 psi; 0.7-1 atm) that was connected to the robot via flexible tubing routed outside of the robot.

#### Gait generation

To model the peristalsis gait and lateral undulation of earthworms, we use the first and last segments as an anchoring/bending segment and middle two segments as an elongation segment. Note that, the middle two segments also have the same structures (with two inner chambers) as the anchoring/bending segments. We used two-chamber elongation segments instead of the one-chamber to eliminate the errors (such as uneven thickness of the elastomeric body) during the fabrication steps which causes varying degrees of undesired deformations under the same pressure. The two-chamber design allows us to adjust the maximum pressure of the segments individually, which provides symmetrical straight elongation.

The gaits were empirically determined by actuating the segments with different sequences. The fastest sequence (with 7±0.6 cm/cycle) we observed was to inflate segments 4-3-2-1 and exhaust them in the same order (Table 1, Gait-D). A cycle starts from the rest state (all the segments are deflated). Pressurization of segment 4 (S4) bends the robot’s back to the left/right (~60*°*) and anchors the body to avoid sliding backward. Then the middle segments (S3 and S2) elongate to provide forward displacement while keeping S4 inflated and finally pressurization of S1 bends the tip of the robot and establishes an anchoring point at the front to propel the back of the robot forward when all the segments (S2, S3, and S4) are deflated. The mean±SD values of the displacements per cycle for other gait sequences are given in Table 1 (SI Movie-4).

**Table 1:** The activation pattern of the segments (Fig. 5a) and displacement of the robot tip per cycle (cm/cyle) for different gait sequences. The robot was tested in a tube (d=4.3 cm). Data shown are means±SD for six cycles/three runs. The shaded row (Gait-D) is the best gait we observed (see SI Movie-4).

| Gait | INFLATE | EXHAUST | cm/cycle |
| --- | --- | --- | --- |
| A | 2-3-4-1 | 2-3-4-1 | $0.4 \pm 0.3$ |
| B | 2-3-4-1 | 4-3-2-1 | $3.3 \pm 0.1$ |
| C | 4-3-2-1 | 2-3-4-1 | $5.1 \pm 0.6$ |
| D | 4-3-2-1 | 4-3-2-1 | $7.0 \pm 0.6$ |

We next demonstrated that without changing the gait sequence, the performance of the locomotion decreases when we do not use undulatory motion. In these experiments, using Gait-D we performed two sets of experiments in a acrylic tube (d = 4.3 cm) and a hexagonal lattice comprised of rigid pegs (cylindrical corks, see Fig. 9) with and without cyclic body undulation. In both environments, the robot that used body undulation (5.94±0.27 cm in the tube, 3.24±0.16 cm in the lattice) outperformed the one without body undulation (2.7±0.16 cm in the tube, 0.65±0.11 cm in the lattice). Fig. 6a shows a gait cycle of an undulated robot when crawling in a tube. Note that, when inflated, the diameter of the segments can increase up to 50% of its neutral diameter, which we used to anchor to the body in the absence of undulation. Data shown in Fig. 6b are the mean±SD for six cycles/three runs with and without body undulation.

**Figure 6:**
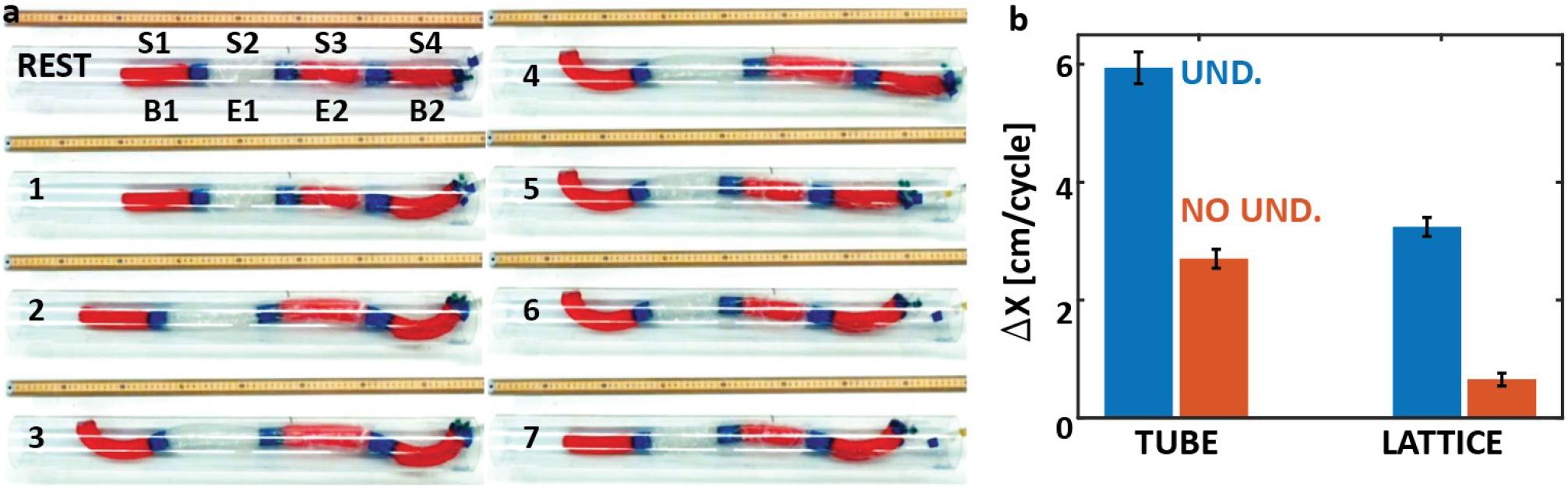
Locomotion performance of the robot with and without body undulation. A gait cycle of an undulated robot. The segments (S1 to S4) were inflated in a sequence of 4-3-2-1 and exhausted in a same order. The front (B1) and back (B2) are the bending segments and middle two segments (E1 and E2) are the longitudinally extending segments. When there is no undulation, the front and back segments only extend. **b.** Displacement per gait cycle with (blue) and without (red) tip undulation in the tube (d = 4.3, level) and lattice (level). Data shown are the mean±SD for six cycles/three runs.

**Figure 9:**
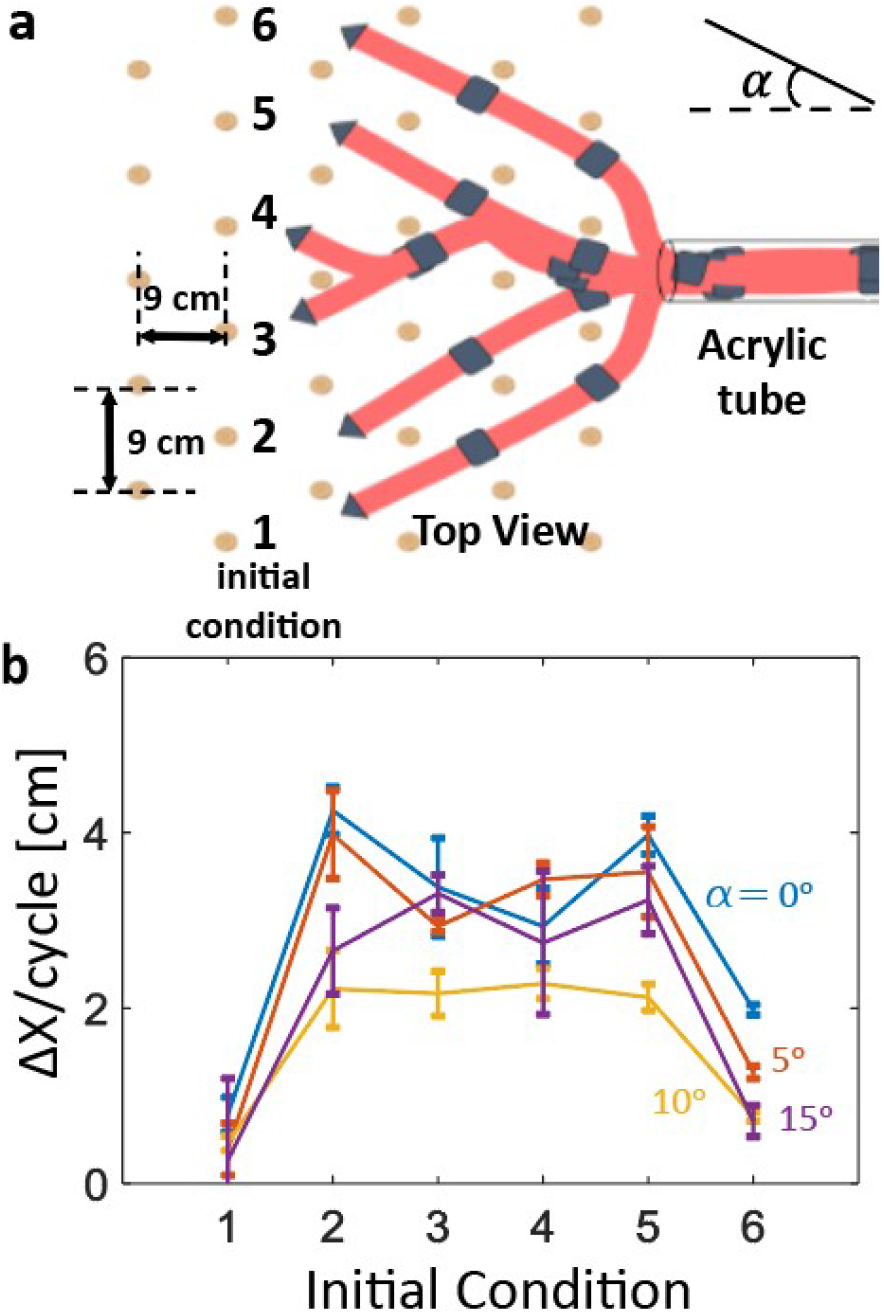
Robot experiments in the heterogeneous environment. (a) Top view of the experimental setup. The cylindrical rigid cork pegs (d=2cm, h=4 cm) were mounted on a melamine laminated board (0.5×1m) with a horizontal and vertical distance equals to 9 cm. An acrylic tube (d= 5.3 cm, l=25 cm) was placed to starting point. The angle of the board (*θ*) is the angle between the horizontal plane and the board surface. The robot was started with six different initial shapes and tip positions. (b) Displacement per cycle as a function of six initial conditions given in (a) for four different board angle (*α* = 0, 5, 10, and 15*°*). Each data point shows mean and std of three experiments.

#### Systematic robot experiments

Our next step is to systematically test the effect of both gait and substrate properties on locomotor performance. During our experiments, we observed that when the friction between the elongation segments and the surface was high, the robot could not overcome friction and would slide back, or undesired body buckling occurred. When we covered the middle two segments with a Polytube, we reduced the friction about 40% and obtained a 40% increase in the tube climbing performance. Therefore, throughout the remainder of the study we used Polytube covered segments.

We first did experiments in four acrylic tubes with diameters *d*_1_=4.3, *d*_2_=5.3, *d*_3_=6.3, *d*_4_=8.3 cm which are 1.7, 2.1, 2.5 and 3.3 times bigger than the robot diameter and changed the tube angle from 0 to 90*°* with an increment of 15*°* (Fig. 7a). In all experiments we used Gait-D that was described in the previous section. Fig. 7b shows snapshots from one of the experiments where the robot climbed in a tube (d=5.4 cm, *α*=90*°*, SI Movie-5). As seen from the results (Fig. 7c), the robot performed similar for all tube angles (5.06±0.95 cm/cycle) until the tube diameter 6.3 cm for all tube angles without any significant reduction in performance. The lateral undulation guarantees that the robot can climb up the tube safely and steadily without slip or falling. However, in the large tube (d=8.3 cm) the mean displacement/cycle decreased from 5.72±0.42 cm to 1.78±0.36 cm as the tube angle was changed from 0 to 45*°*. After 45*°* the robot was unable to apply enough lateral force that required to overcome the robot weight (200 gr.) during the climbing process and it slid back.

**Figure 7:**
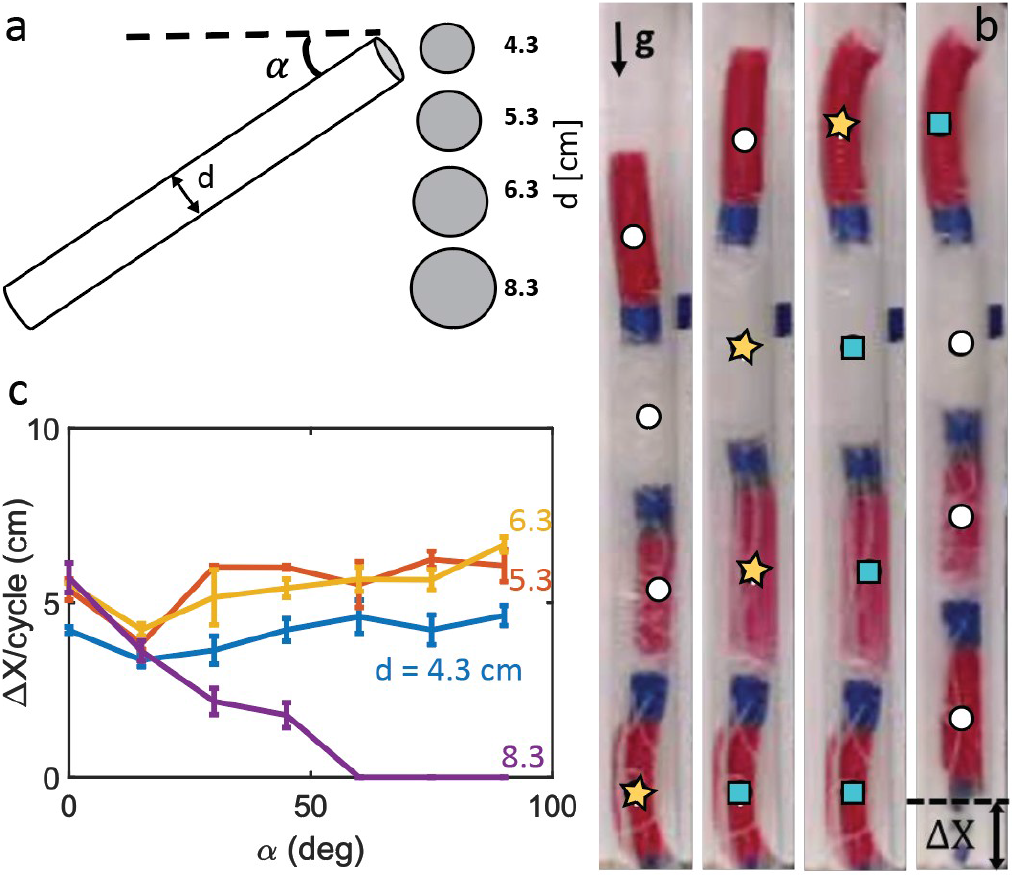
Demonstration of the basic usage of robot body bending as an anchor mechanism to locomote in an acrylic tube. (a) Experimental setup. Four different diameter of tubes (4.3, 5.3, 6.3, 8.3 cm and 1m) are attached to tripod and we changed the tube angle (*α* between horizontal plane). (b) Example climbing test in a 90*°*, d= 5.3 cm tube. Colored shapes represent actuation state of the segment (yellow star: actuate, blue rectangle: actuated from previous step, white circle: unactuated). The longitudinally extending segments (2^nd^ and 3^rd^ segments) were covered with PolyTube to reduce the friction. (c) Displacement per gait cycle (Δ*X*) as a function of *α* for different diameter tubes (blue= 4.3, red= 5.3, yellow= 6.3, green= 8.3 cm).

### 2.3. Heterogeneous space animal experiments (undulation aided grabbing)

In previous sections, we discovered that lateral undulation conferred benefits to the worm’s subterranean movement in the natural and laboratory confined environment. Given these locomotion benefits we next examined how the worms actively control the body shape when they encounter obstacles. As a model of surface irregularity, we studied worms as they crawl through an inclined (25*°*) obstacle course (l = 40, d = 24 cm) made up of regularly distributed rigid pegs (h = 1.5 cm, d = 3 mm, Fig. 8a). To encourage worms to use pegs instead of surface friction, we covered the surface of the test area with a low friction material (Teflon sheet). Fig. 7a shows one of the example combined time-lapse images of a worm while climbing over the obstacle course (SI Movie-3). Rather than continuing to travel through the incline in a straight line, they searched the environment, grabbed the pegs by bending the body (especially tip) and used pegs as anchor points to assist their movement (8 animals, 3 trials each).

**Figure 8:**
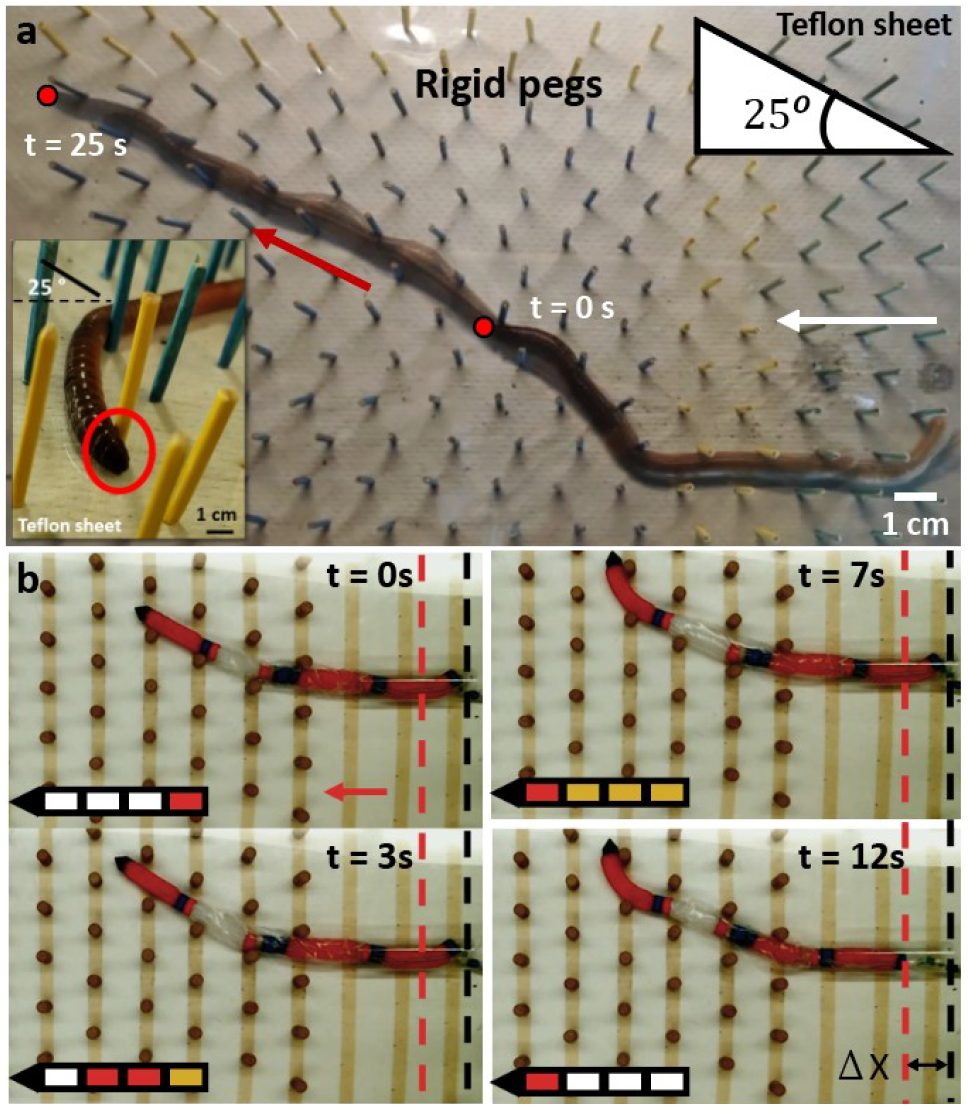
The usage of undulation for hooking and anchoring in heterogeneous environment. (a) Earthworm is climbing on a slippery inclined surface. The surface of the incline (25*°*) was covered with a Teflon sheet to prevent the worm from benefiting from surface friction and encourage the use of rigid pegs. (b) Snapshots from one of the robot experiment. The robot started from the initial condition five (Fig. 9a) and locomoted to the left (red arrow) using the Gait-D given in Table 1. Insets show the actuated segments (red: actuated, yellow: actuated from previous step, white: unactuated)

### 2.4. Heterogeneous space robot experiments

To show that the robot benefits from lateral undulation similar to animal we next performed experiments in a heterogeneous environment (SI Movie-5). We mounted cylindrical rigid cork pegs on a melamine laminated board in the form of a vertical hexagonal lattice Fig. 8b. To provide initial anchoring point at the back we used an acrylic tube at the middle of the setup whose length equals to approximately one segment length and started the robot from six different initial configuration as shown in Fig. 9a. We also changed the angle of the surface (*α*) with respect to the ground from 0 to 15*°* with an increment of 5*°* to demonstrate that lateral undulation is beneficial in inclined heterogeneous environment. Using Gait-D, we performed three experiments in each an initial condition and measured the displacement per cycle. Fig. 8b shows snapshots from one of the experiment (*α*=0*°*) where the robot was locomoting within the obstacles and using pegs for holding points. The robot performed similar (2.33±1.25 cm/cycle) for all the surface angles (0 to 15 *°*) when it started from the initial conditions two to five (Fig. 9b). The open loop tip bending allowed the robot to overcome the obstacles without any sensory feedback and complex control methods. However, if the relative angle between segments three and four was bigger than 45*°*, which happened when the robot started from initial condition 1 and 6, the buckling of the extending segments prevented the robot moving forward.

## 3. Conclusions

In this paper, we studied lateral undulation of earthworms, which, to the best of our knowledge, has not been studied before. We showed with systematic laboratory experiments that lateral undulation helps to keep the animal from slipping while climbing in confined environments (different size inclines (0 to 90*°*)) such as under soil tunnels and utilizes locomotion in heterogeneous environment.

Soft robots possess the benefits of having flexible bodies that promote robustness during locomotion while maintaining animal-like motion characteristics. To test our biological observations, we built a robophysical model of an earthworm which has a segmented soft body that can laterally bend and elongate. We show that an open loop body undulation strategy can be used to propel the robot in complex environments. The robot can climb in tubes that are more than three times larger than its diameter and navigate through obstacles without any external or internal sensing.

For advanced applications of autonomous robots, such as search-and-rescue operations and sub-surface soil exploration and monitoring, robots have to move through rough terrain with an ability to control its deformation and sense the environment [52]. Our recent study has shown that the soft, stretchable nanocomposite strain sensors integrated with the robot body can be used for feedback-controlled locomotion in complex and dynamic terrestrial environments [53].

## Acknowledgment

This material is based upon work partially supported by the National Science Foundation under Grant No. 1545287. Funding for Y.O.A and D.I.G provided by ARO and ARL MAST CTA; funding also provided by Dunn Family Professorship. P.L. supported by NSF Grants 1908042, 1806580, 1550461. Any opinions, findings, and conclusions or recommendations expressed in this material are those of the author(s) and do not necessarily reflect the views of the National Science Foundation. We also would like to thank the Georgia Tech IRIM Seed Grant program for supporting this research.

## Author contributions

Y.O.A and B.L fabricated the robot; Y.O.A. and A.C.F. performed animal experiments, Y.O.A designed and performed the robot experiments; M.S. aided in robot experiments; Y.O.A., F.L.H. and D.I.G. wrote the paper; F.L.H. and D.I.G. guided the overall research program.

